# Manipulation of rhizosphere microbiome by *Microbacterium* sp. GB16_1_BI to promote plant growth

**DOI:** 10.64898/2026.05.15.725310

**Authors:** Papri Nag, Rajani Govindannagari, Kavuru Prasad, Tummala Mounika, Latha P. Chandran, Sampa Das, Prasad Babu MBB, Sundaram RM

## Abstract

Plant growth promoting microbes enhance developmental progression of the host by influencing its nutrient availability or by deploying secondary metabolites responsible for manipulating the hormonal crosstalk. *Microbacterium bengalense* sp. nov. GB16_1_BI (Accession number: SRX9280401), a newly identified ammonium releasing Actinomycetota, could enhance plant growth by manipulating rhizosphere bacteria. Amplicon sequencing of the 16S rRNA V3-V4 region from the rhizosphere of the black rice (Chakhao Poireiton) showed that GB16_1_BI could inhibit most bacteria. However, GB16_1_BI inoculation encouraged the growth of rare bacteria specific to waterlogged rice rhizosphere. Analysis of the OTUs using PICRUSt2 (Phylogenetic investigation of communities by reconstruction of unobserved states) showed increased abundance in the marker genes for nitrogen cycling (*nifH, nrfA* and *nrt*) but not for *nifD* or *nifK* which was also reflected in the ANOSIM analysis in the OTUs of the N-fixing bacteria. Marker genes for methane metabolism (*comA, comB, cofG* and *cofH*) were also more abundant in the inoculated plants than the control; however, ANOSIM studies did not support this observation in the OTUs of methane cycling bacteria. Both *Methylosinus* and *Methylocystis*, the two most abundant methanotrophic OTUs, are also known to be nitrogen fixers. Hence, GB16_1_BI could influence plant growth predominantly by manipulating nitrogen cycling microbes. The genome sequence as well as untargeted metabolome analyses of GB16_1_BI showed abundance of secondary metabolites with probable antimicrobial activity. GB16_1_BI could utilize varied carbohydrates and amino acid as energy source and form persister-like cells may help it to survive in the soil in absence of the host plant.

## Introduction

Since ancient times rice has been cultivated in the river valleys (Civáň et al., 2015). These soils were usually fertile due to the floods bringing in the humus and topsoil from the surrounding upland areas. Expansion of rice cultivating areas due to population increase led to an escalation in irrigation and synthetic fertilizers use. However, low utilization capacity of the rice plants for the applied nitrogenous fertilizers resulted in land and water pollution (Erisman et al., 2013; Zhang et al., 2015). This has necessitated the use of sustainable methods of supplying nitrogen to the plants (Coggins et al., 2025). Traditional rice cultivation methods require large amounts of water and have the inherent problem of methane production (Ehhalt and Schmidt, 1978; Nouchi et al., 1990; Bhattacharyya et al., 2019). In recent years, to mitigate the effect of water requirement and methane production, agronomic practises such as drip irrigation, alternate wetting and drying cycles and direct seeded or upland cultivation practices are being followed (Matsuda et al., 2022).However, alternative wetting and drying, and upland practises are prone to another form of pollution: production of nitrous oxide, also a potent greenhouse gas (GHG) (Kritee et al., 2018).Methanogenesis, nitrous oxide production and soil nutrient availability are largely microbial processes and can be managed by sustainable methods like rhizosphere engineering.

Soil contains innumerable species of microbes, while almost all contribute to the biogeochemical cycles, not all are beneficial for growth of crop plants. Microbes can compete with the plants for nutrients (Klironomos, 2002; Fitzpatrick et al., 2018) and contribute to harmful GHG emissions (Kirschke et al., 2013; Lagomarsino et al., 2016). Rhizosphere engineering can provide only short-term benefits by introduction of beneficial microbes for mitigating these harmful effects. The soil microbiome is considered a resilient community of microbes and external application of high density of inoculants are outcompeted by the native microbes (Wang et al., 2021). Establishing a new strain of bacteria can produce a shift, albeit transiently, in the niche structure of the native microbes (Mallon et al., 2018). Inclusion of keystone species (Carlström et al., 2019; Laurich et al., 2024) or those which can interact with native keystone species may have a stronger effect on shaping the microbiome (Sun et al., 2025). In fact, native microbes are the most important factor in assembly of the rhizosphere microbiome with the root exudate being responsible for enrichment of specific microbes (Bulgarelli et al., 2013). In rice, different growth stages involve varied water requirements thereby forming soil conditions which encourages the growth of microbial consortia with differential oxygen requirements (Revsbech et al., 1999). The oxic-anoxic condition encourages the growth of methanogens (generating methane) and methanotrophs (utilizing methane) (Lee et al., 2014; Shen et al., 2014; Ding et al., 2019). Methanogens are mostly anaerobic Archaea (Baun, 2018) while methanotrophs can be both aerobic and anaerobic (Guerrero-Cruz et al., 2021). Rice roots contain a special tissue, aerenchyma, responsible for oxygen supply to the roots and release of methane to the environment (Nouchi et al., 1990). Aerobic, facultative aerobic and microaerobic microorganisms can survive near the roots and endophytic regions due to the oxygen supplied by the aerenchyma tissues (Bao et al., 2014). The isolation of a novel species of *Microbacterium* designated as *M. bengalense* GB16_1_BI sp. nov. (Accession number: SRX9280401) which can secrete ammonium has been reported previously (Nag et al., 2026). This study demonstrates the potential of GB16_1_BI as a plant growth promoting bioinoculant and its effect on the rhizosphere microbiome of rice.

## Methods

Bacterial growth conditions: GB16_1_BI was maintained on N-free plates at 28 °C. Liquid cultures were always maintained under shaking. For inoculation in plant growth assays, the N-free liquid cultures were supplemented with 0.25 gm l^-1^ yeast extract and grown for 7-10 days for depletion of the nitrogen in the media.

Whole genome sequencing: Genomic DNA isolation was done from cells cultivated overnight in N+ medium (N-free medium + 0.25 gm l^-1^ yeast extract). Library preparation and sequencing was done by Genotypic Technologies Ltd (Bengaluru, India). In brief, the native barcoded library preparation was cleaned, and the library was quantified on Qubit 4 Fluorometer. Sequencing was done on Nanopore PromethION sequencer, (PromethION24 and Data Acquisition Unit, Oxford Nanopore Technology, UK) using PromethION flow cell (FLO-PRO114M). Porechop-v0.2.4 (Wick et al., 2017) (https://github.com/rrwick/Porechop) was used for removal of adapter sequences. The nanopore processed reads were used for de novo assembly using Flye-v2.9.5-b1801 assembler (Kolmogorov et al., 2019). The primary read statistics was generated using NanoStat-v1.6.0 (De Coster et al., 2018). The quality of the genome sequence was assessed using gVolante (Nishimura et al., 2017) for completeness. The genome was annotated using RAST (Brettin et al., 2015), Prokka [v.1.12] (Seemann 2014), Pfam (Mistry et al., 2021), NCBI (Tatusova et al., 2016) and KEGG (Aramaki et al., 2019).

For analysis of carbohydrate utilization KEGG (Kanehisa et al., 2020), dbCAN2 (Zhang et al., 2018) and CAZy (Lombard et al., 2013) was utilized. For identifying genes functioning in DNRA pathway, Pfam (Mistry et al., 2021) sequence search, NCBI CDD BLAST (Mount, 2007), Kofam (Aramaki et al., 2020), KEGG mapper (Kanehisa et al., 2020) were used. AntiMASH was used for predicting regions of biosynthetic clusters (Blin et al., 2023).

Biochemical assays: Ammonium estimation was done by Nessler’s reagent (Paustian et al.,1989). Nitrate and nitrite estimation was done by Griess reagent (Syre et al., 2003). Protein estimation was done by Bradford’s method. Briefly, nitrate, nitrite, ammonium and protein estimation was done from each stage with a starting cfu count of 10^6^, under shaking @ of approximately 100 rpm with temperature maintained at 28 °C.

Transcriptome sequencing: RNA extraction from GB16_1_BI was done using Trizol reagent from log-phase cells. RNA Seq analysis was done using Illumina NovaSeq6000 platform (Sandor speciality diagnostics, Hyderabad, India). Comparative analysis was done after normalization. Annotation of the identified sequences were done using BLAST from NCBI, and Uniprot database (The UniProt Consortium, 2009).

LCMS/MS analysis: GB16_BI was grown in LB and LBS medium for 72 hours and the whole cell extracts (French press) were run on 12% SDS PAGE gel, bands were excised and subjected to LC-MS/MS after trypsinization (Gundry et al., 2010) in a Waters, XeVo G2 XS QTof (Bose Institute, Kolkata, India). Using ProteinLynx Global Server version 3.0.3 (Waters, Miliford, USA), peptides were identified. The peptides were annotated using the UniProt database (The UniProt Consortium, 2009).

Extraction of metabolite was done with 80% ethanol from log phase GB16_1_BI cell pellet followed by filtering with 0.22um filter. LCMS (Shimadzu Prominence-I LC interfaced to a Shimadzu triple quadrupole LCMS-8045 mass spectrometer; Shimadzu Corp., Kyoto, Japan) was used for electrospray ionization (ESI)–MS analysis and full-scan mass-spectra were acquired over a mass range of m/z 50–2000. MZmine was used for mass feature detection and identification. with the MZML raw LC/MS files cropped to a retention time range of 0–45 min. Biocyc/Pubchem databases were used for metabolite identification (Nag et al., 2026).

Plant growth conditions: Chakhao Poireiton, a black rice variety was selected for the plant assays. Plants were grown under greenhouse conditions in non-sterile, soil from rice field (Madhyamgram field station of Bose Institute, West Bengal, India) and mixed with dried FYM thoroughly to ensure similar composition in earthen pots. Plants were grown without the application of synthetic fertilizer. Greenhouse temperature was maintained at 34-36 °C and humidity at 70-80%. Light conditions were kept at 8 hours light and 16 hours dark. Seeds were surface sterilized using sodium hypochlorite followed by 70% ethanol and washed properly with autoclaved water before sowing on autoclaved agro peat. Seedlings were transplanted 10 days after germination and observations were recorded 30 days after transplanting.

For analysis of root microbiome, plants were grown in 1m^2^*0.5m^2^ pots in three replicates (each pot had N=6 plants) during rice growing season in an open area under natural humidity, temperature and daylight conditions during the Rabi season (January-March 2025). The nursery was maintained on soil (ICAR-IIRR fields) without application of nitrogen and 30-day old seedlings were transplanted in 1m^2^*0.5m^2^ pots with standing water. The pots were always maintained under waterlogged conditions with borewell water. Sample collection for soil DNA isolation was done 60 days after sowing and the soil was stored at -80 °C. Plant tissue samples were frozen immediately in liquid nitrogen and stored at -80 °C until RNA extraction and cDNA preparation.

Plant inoculation assay: For plant RNA extraction and CHN analysis, plants were inoculated with live GB16_BI (LGB) and heat killed GB16_1_BI (HKGB). GB16_1_BI was cultivated in N-free medium for 72 hours and harvested by centrifugation @ 8000 rpm. The pellet was washed twice with 1/4th volume of initial culture and diluted @ 10-8 ml^-1^ in autoclaved water and divided into two parts. One portion was used for producing HKGB inoculant preparation by autoclaving. Root-dip treatment was done for 10 minutes in HKGB and LGB, before transplantation into pots.

Rhizosphere microbiome analysis was done form plants maintained in pots under natural conditions. GB16_1_BI cells were cultured in N-limited medium (modified Burk’s medium containing 0.25 gm l^-1^ yeast extract for 7-10 days and diluted to 10-8 cell ml^-1^ with water. Root-dip treatment was extended to 2 hours for the microbiome assay.

CHN analysis: CHN analysis was done at the SAIF facility of Indian Institute of Technology, Bombay using the CHN(O) analyzer (ThermoFischer). All experiments were done with three biological replicates.

Plant RNA extraction and qRT-PCR analysis: RNA was extracted from the frozen plant tissues harvested 30 days after inoculation. The tissues were ground on a mortar-pestle with liquid nitrogen for Trizol (Invitrogen) extraction as per manufacturer’s instructions. The isolated RNA was treated with DNase (Fermentas) to remove any residual DNA. First strand cDNAs were synthesized using High-capacity cDNA reverse transcription kit (Applied Biosystems, ThermoFisher). qRT-PCR was performed using Applied Biosystems™ Power SYBR™ Green PCR Master Mix (Thermo Fisher Scientific™) on the Applied Biosystems 7500 fast real-time PCR machine.

Amplicon sequencing for 16S rRNA V3-V4 region: Rhizosphere soil along with the root was macerated for soil DNA extraction using the PowerSoil DNA isolation kit (Qiagen) and quantified on nanodrop. 16S rRNA V3-V4 metagenomic sequencing and downstream analysis was outsourced for MiSeq (Illumina Inc). The 16S rRNA V3-V4 amplicon libraries were constructed in alignment with the library preparation protocol from Illumina Inc. Briefly, 12.5 ng DNA was amplified using16S rRNA V3-V4 primers, the PCR products were bead purified and subjected to another round of PCR with dual indices and adapters to generate the respective libraries as recommended by Illumina, Inc. The cleaned libraries were quantitated on Qubit® fluorometer and appropriate dilutions loaded on a D1000 screen tape to determine the average library size. Raw reads from the MiSeq sequencing system were pre-processed by FASTP (Chen et al., 2019) for achieving <Q20 score. Pair end sequences were assembled using PEAR (Zhang et al., 2013) and low-quality reads were filtered out. Taxonomic profiling was done using the Kraken2 database (Wood et al, 2019). Phylogenetic investigation of communities by reconstruction of unobserved states was done using the PICRUSt v2-2.6.3 (Douglas et al., 2020).

## Results

Genome analysis of GB16_1_BI: Sequencing of GB16_1_BI at a depth of 350X provided a better understanding of the physiology of the isolate. A total of ∼1.42 GB nanopore long reads data was generated corresponding to 350X coverage. *De novo* genome assembly of the bacterial sample resulted in single contig with draft genome of size ∼3.59 MB and N50 value of ∼3.59 MB. The quality of the genome sequence was assessed using gVolante (Nishimura et al., 2017) which uses BUSCO scores showing 99.2% completeness. The genome was found to be 3,596,333bp with a GC content of 70.7. The genome resolved into 3432 protein cds, 51 RNA and 144 tRNA coding sequences. KEGG, Pfam and PDB analysis revealed versatile nitrogen metabolism pathways. One of the most prominent features was the presence of an intact prophage region identified by PHASTEST (Wishart et al., 2023). The prophage region contained the bacterial attachment region, integrase, head-protein and tail-protein. AntiMASH (version:7.0) predicted seven regions in the genome with biosynthetic gene clusters (BGC). Seven regions containing BGCs with genes expressing RRE-containing (post-translationally modified peptide RiPPs), lanthipeptide class-iii, T3PKS, NAPAA, terpene, lanthipeptide class-i, lipopolysaccharide and beta-lactone. The NAPAA containing cluster showed a 100% match with ε-Poly-L-lysine, a non-ribosomal peptide synthetase of *Epichloe festucae*.

The genome sequence confirmed previously reported complete assimilatory nitrate reduction (ANRA) pathway, an incomplete dissimilatory nitrate reduction (DNRA) pathway with a single component missing (Nag et al., 2026) and the absence of known nitrogen-fixing pathways. The dbCAN2 tool identified 132 active carbohydrate enzymes compared to the earlier reported 117 carbohydrate metabolism genes from the genome sequence of GB16_1_BI using KEGG (Nag et al., 2026).

Quantification of ammonium release: Biochemical assays with Nessler’s reagent showed that GB16_1_BI could release 36 mM NH3 mg^-1^ protein when NO2-was used as the nitrogen source and 126 mM NH3 mg^-1^ protein when NO3-was used (Fig. 2). However, NO2-becomes toxic and no cell growth can be seen above 2mM concentration and NO3-became toxic in concentrations above 200mM. GB16_1_BI released 46 mM NH3 mg^-1^ protein into the medium when no nitrogen source was provided. It may be speculated that GB16_1_BI can survive on very less amount of N by scavenging the NO2- and NO3-formed spontaneously at the water-air interface (Kumar et al., Bose et al., 2024). Cells grown in N-free sucrose medium were reminiscent of the persister cells produced by most bacteria (Fig. 3a, left panel). Persisters are slow-growing or growth-arrested cells, formed by most bacteria to survive stress conditions (Fisher et al., 2017). These persister-like cells continued to release ammonium and produce copious amounts of exudates until sucrose was exhausted from the medium. The recalcitrant nature of GB16_1_BI grown in N-free medium proved to be a deterrent in identifying the pathway for this phenomenon and needs to be explored further. Among other pathways which can produce ammonium as a by-product, the complete pathway for ornithine-ammonia cycle and nitroalkane degrading pathways has also been predicted in the genome of GB16_1_BI.

**Fig. 1.**
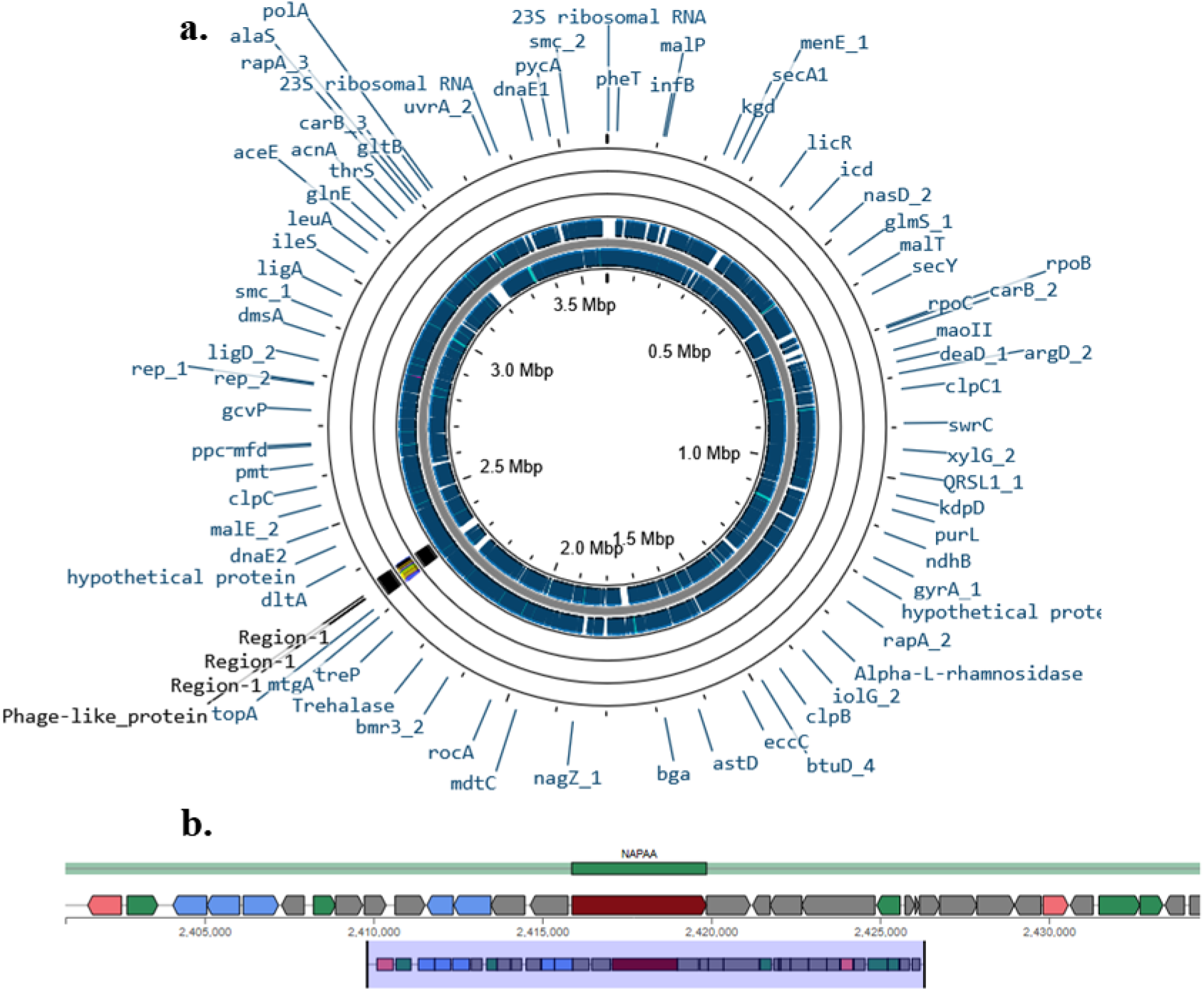
Genome analysis of GB16_1_BI. (a) Schematic diagram of the whole genome sequence annotated by Prokka. **(b)** Secondary metabolite biosynthetic gene cluster predicted by AntiMASH.

**Fig. 2:**
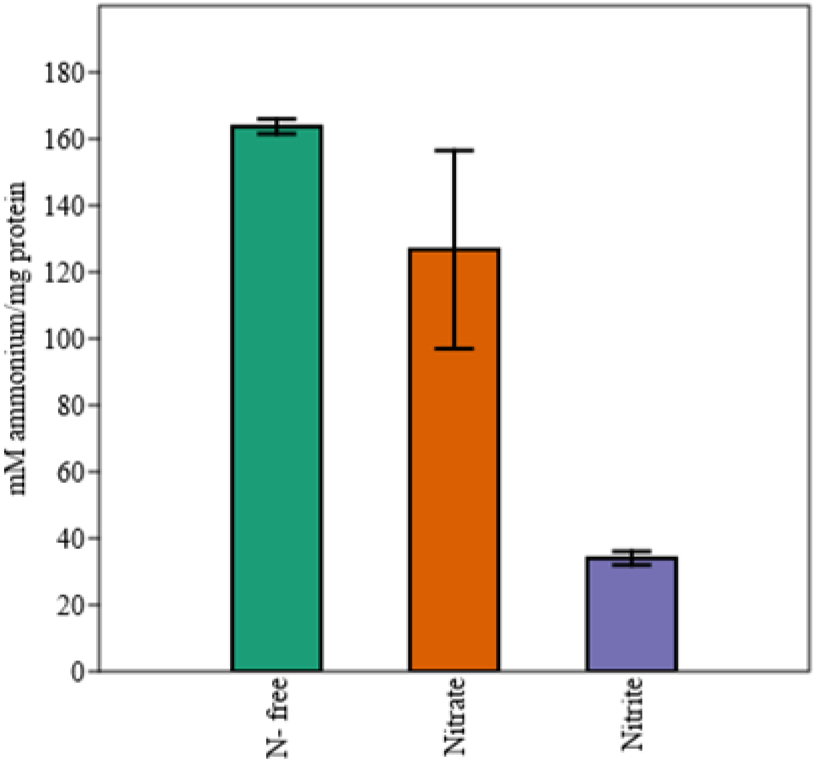
Ammonium released by GB16_1_BI in N-free, nitrate (60 mM) and nitrite (0.8 mM) containing media. The quantification is based on the observations taken at log phase (48 hours) of growth standard error bars represent mM ammonium mg^-1^ protein during these three stages. All experiments were done in triplicate.

**Fig. 3.**
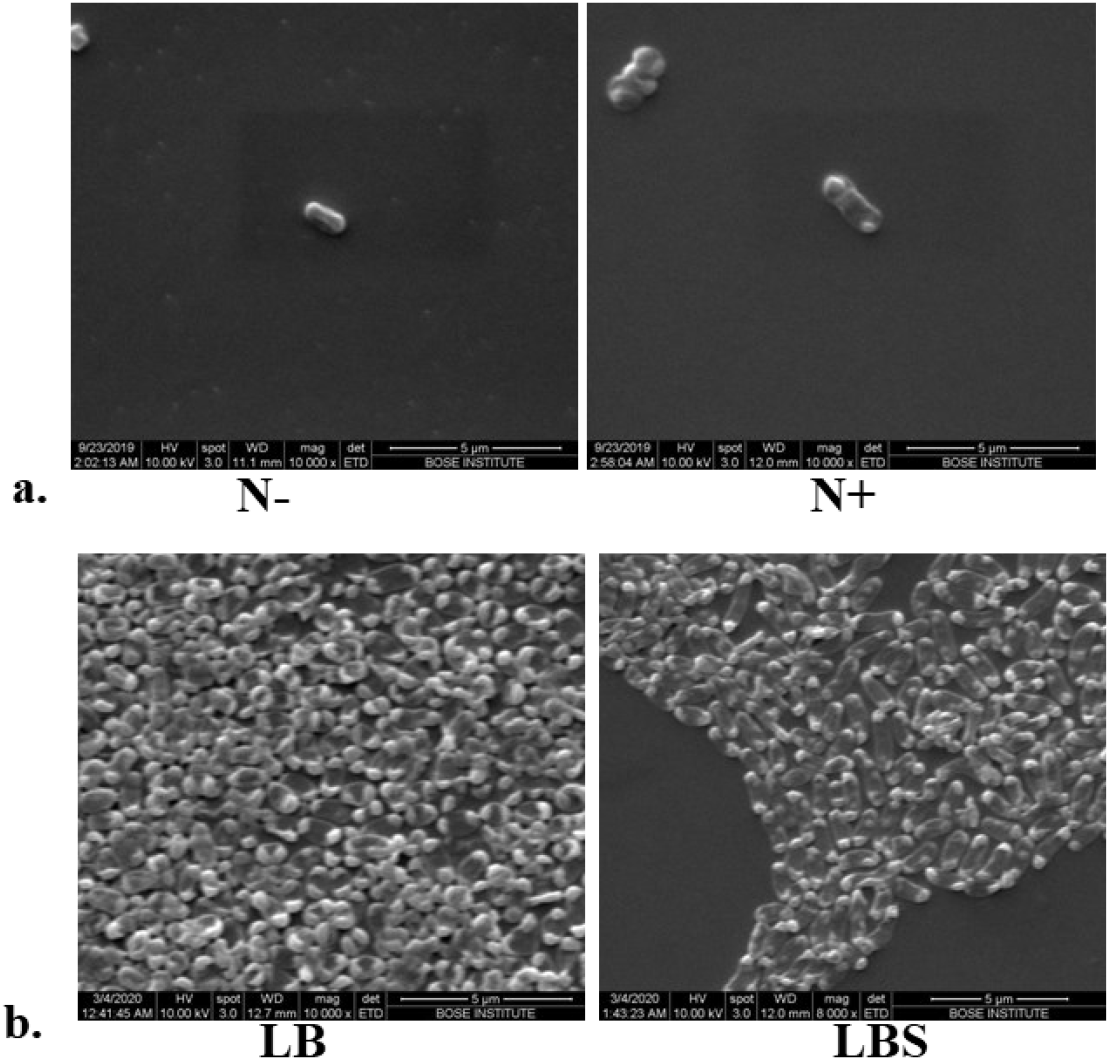
Scanning electron microscopic examination of GB16_1_BI under different nutritional conditions. **A**. cells grown in N-free and N+ medium. **B**. Cells grown in LB and LBS (LB+Sucrose) medium.

Analysis of carbohydrate utilization: GB16_1_BI had the ability to utilize diverse carbohydrates as energy source (Nag et al., 2026). Using the Ecoplate (BIOLOG Inc., USA) assay it was observed that GB16_1_BI could utilize almost all the carbon sources provided except, xylose, tween 40 and tween 80. Sucrose was selected for all further studies as the lag phase was shortest in sucrose as judged by OD at 600nm. Agar plates containing sucrose were distinguished by the excessive production of exudates on both N-free and N-containing (LB) media. The morphology of GB16_1_BI cells was elongated in LB+20% sucrose (LBS) with faster growth (Fig. 3b).

To understand the effect of sucrose on cell growth, transcriptome analyses and changes in protein profiles of cells grown in LB and LBS broth were done. Transcriptome analysis showed upregulation of several genes encoding for sugar transport and assimilation in LBS. Genes for growth, cell division, DNA helicases and elongation factors were also upregulated in LBS. The upregulation of transcriptional regulators identical to sporulation factor regulators in LBS indicated a modulation of lifestyle in GB16_1_BI in presence of sucrose. Among the highest upregulated transcripts, VOC related protein, transferase and antibiotic monooxygenase, important traits for survival, were upregulated in LBS. The secretory proteins SecA and SecY were also upregulated in LBS. Protein profile analyses of bands excised from 1D PAGE gels visibly different in LB and LBS were used for LC-MS/MS analysis. Both the protein profile and transcriptome analyses revealed upregulation of amino acid dehydrogenases and glutamine synthetase (GS) in LBS. In addition, a high mobility group (HMG) protein identified as Histone binding (HU) protein was expressed in LBS.

Metabolite production by GB16_1_BI: Secondary metabolites produced by GB16_1_BI was identified using untargeted metabolite analysis in positive and negative scan mode of the Shimadzu LCMS-8045 using the MZmine software. Most significant observation from the whole cell metabolite analysis of GB16_1_BI was the identification of metabolites with antimicrobial signatures. Concurring with the genome sequence data, untargeted metabolite analysis of GB16_1_BI cells identified the production of phenazines.

The parent pathways responsible for production of 47%, 22% and 14% of the total secondary metabolites (SM) were the amino acid, the polyketide and the alkaloid pathways respectively (Fig. 4a). Among the 20 most abundant SMs, analysis for clear peak and retention time identified signatures for two antimicrobials and metabolites which are known to modulate cell wall (Fig. 4b). Concurring with the AntiMASH data two catechols-Resorcinol and alkylresorcinol and phenazine derivatives-methyl phenazine-1-carboxylate and 1,6-phenazinedimethanol were identified.

**Fig. 4.**
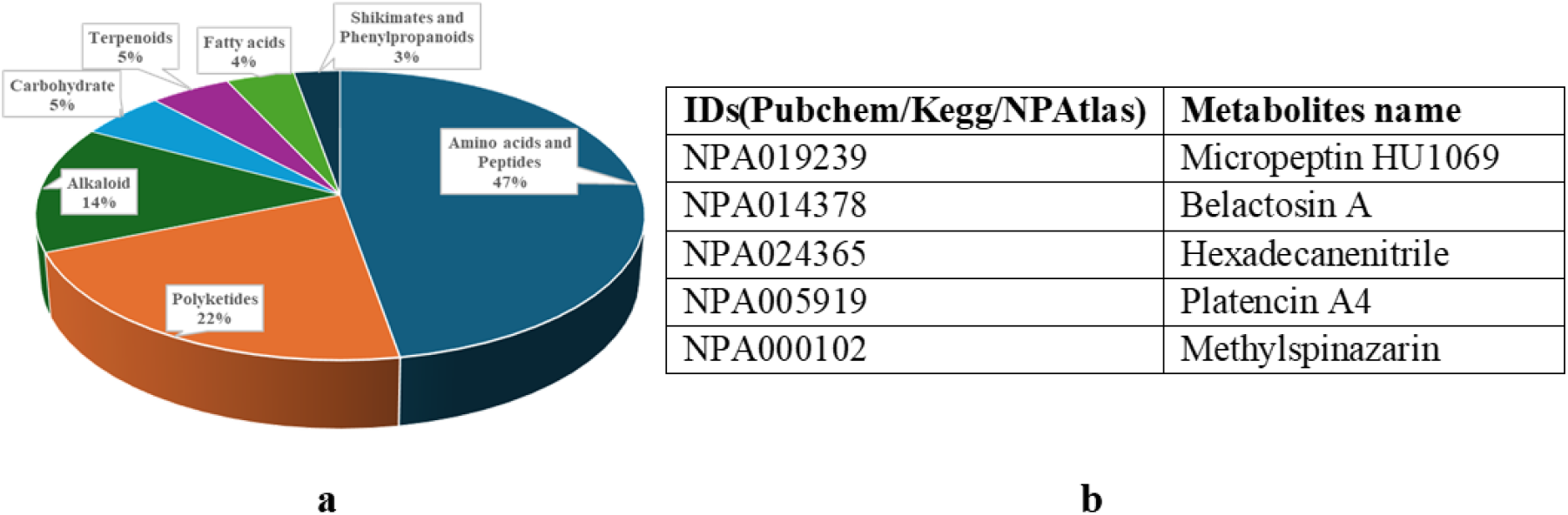
Classification of metabolite produced by GB16_1_BI on LB medium based on **(a)** natural product classifier pathways. **(b)** the highest peak area (five most abundant metabolites are shown)

Plant growth promotion studies: Root dip inoculations with GB16_BI significantly increase the height and fresh weight of the plants compared to uninoculated controls (Fig. 5). Bacterial count in potted soil after one month of transplantation showed that GB16_1_BI could be detected in the root endophytic regions, although not all roots harboured the endophyte.

**Fig. 5.**
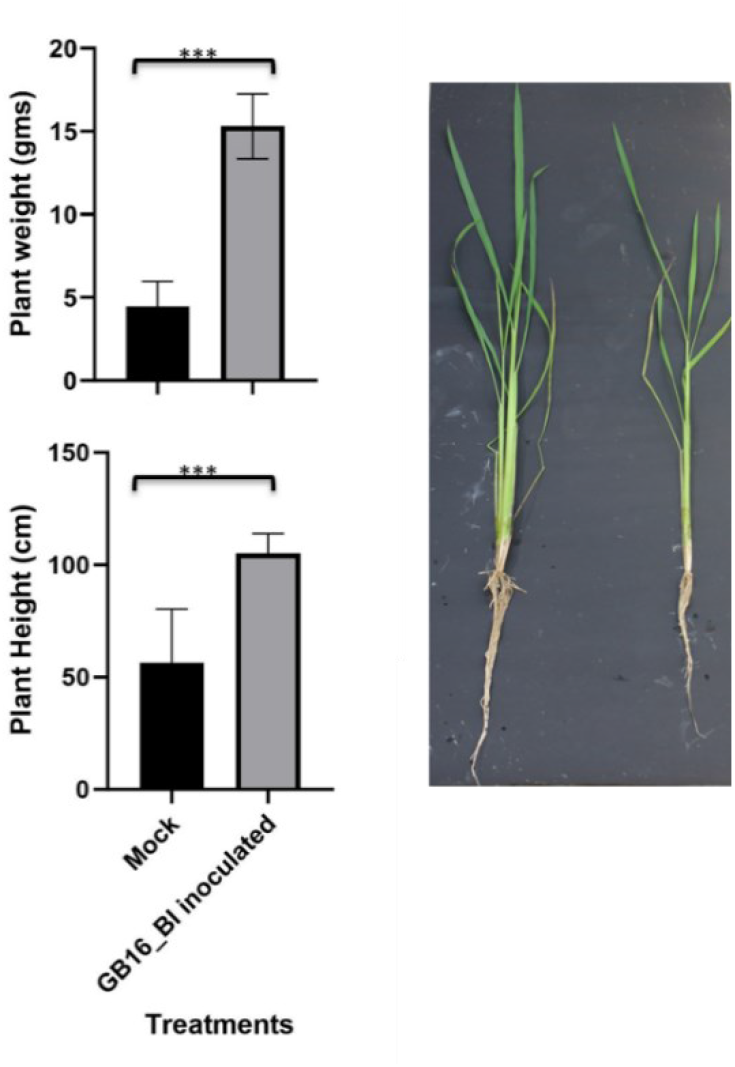
Plants inoculated with GB16_1_BI are taller and have higher fresh weight compared to mock inoculated plants. Values are the mean ±sd (n = 10). Significant differences from WT were determined by Student’s t tests; ***P< 0.005.

To ascertain that the carried over nutrients present in the inoculation medium did not affect gene expression studies and the CHN analysis, control plants were inoculated with heat killed GB16_1_BI (HKGB).

Elemental analysis for carbon, hydrogen and nitrogen (CHN) in plant tissues showed that LGB inoculated plants had higher N content than HKGB plants, although the difference was not statistically significant. It must be noted here that HKGB (due to release of cell content after heat killing) also acted as a source of nitrogen, thus diminishing the effect of N released by live GB16_1_BI (LGB). However, the carbon and hydrogen content was significantly elevated in plants inoculated with LGB compared to the HKGB (Fig. 6a).

**Fig. 6.**
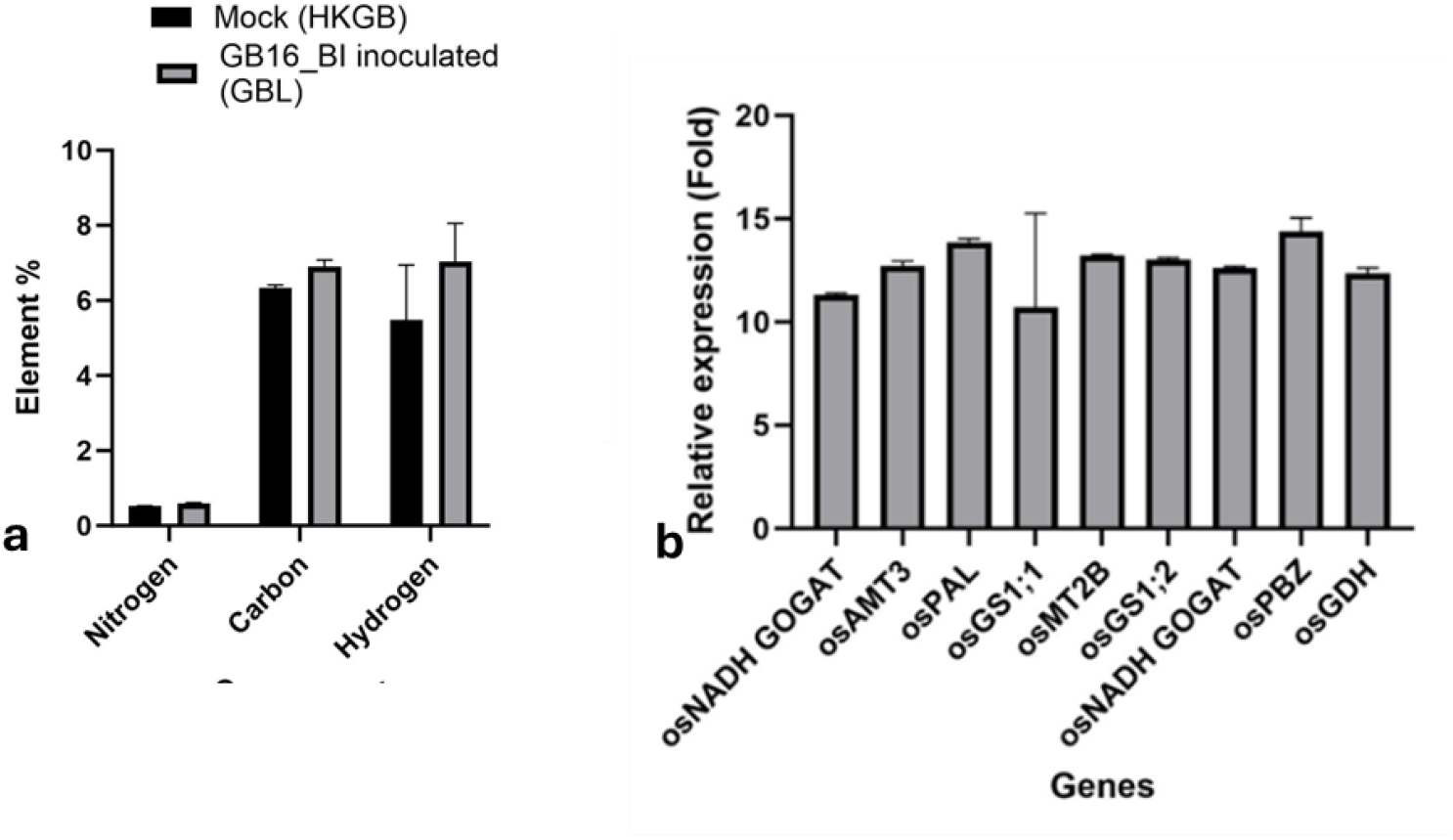
(a) Element (CHN) analysis of plants inoculated with live GB16_1_BI (n=3). Values are the mean ±SD (n = 6, P= ns, not significant). (b) The expression of osGS1;2 and osGOGAT is higher in LGB plants compared to HKGB plants. Values are the mean ±SD (n = 6, P< 0.005).

Expression analysis of genes which are known to be influenced by the N status of the plant growth medium was done to understand the effect of NH3 released by GB16_1_BI. AMT3 coding for ammonium transporter; GS1;1 & 1;2 coding for Glutamine synthetase; NADH GOGAT 1 & 2coding for Glutamate synthase was upregulated in plants inoculated with LGB compared to HKGB inoculated plants (Fig. 6b). Glutamine synthetase (GS) was expressed approximately 3-fold higher in LGB plants compared to HKGB inoculated plants. Microbes (both beneficial and harmful) present in the rhizosphere and phyllosphere can modulate the immune pathway of host plant (Zhang, 2023). Three genes which are considered important for modulating the immune response in rice during microbiome-plant interactions: Phenylalanine ammonia-lyase, Metallothionein-like protein 2C and PBZ1 pathogenesis-related Bet v I family protein was upregulated in LGB plants compared to HKGB inoculated plants (Fig. 6b).

Microbiome studies: To understand the effect of LGB inoculation on rhizosphere microbiome, 16S rRNA amplicon sequencing was done after 30 days of transplantation. MiSeq analysis identified 3969 distinct OTUs belonging to 51 phyla (Fig. 7a) and 3649 genera. Fold change analysis showed 41% of OTUs from LGB plants were lower in abundance than the control plants, 12% had higher fold-change and only 2% of the OTUs showed no fold change due to GB16_1_BI inoculation. Rare OTUs were highest in proportion in both uninoculated control and LGB plants.

**Fig. 7.**
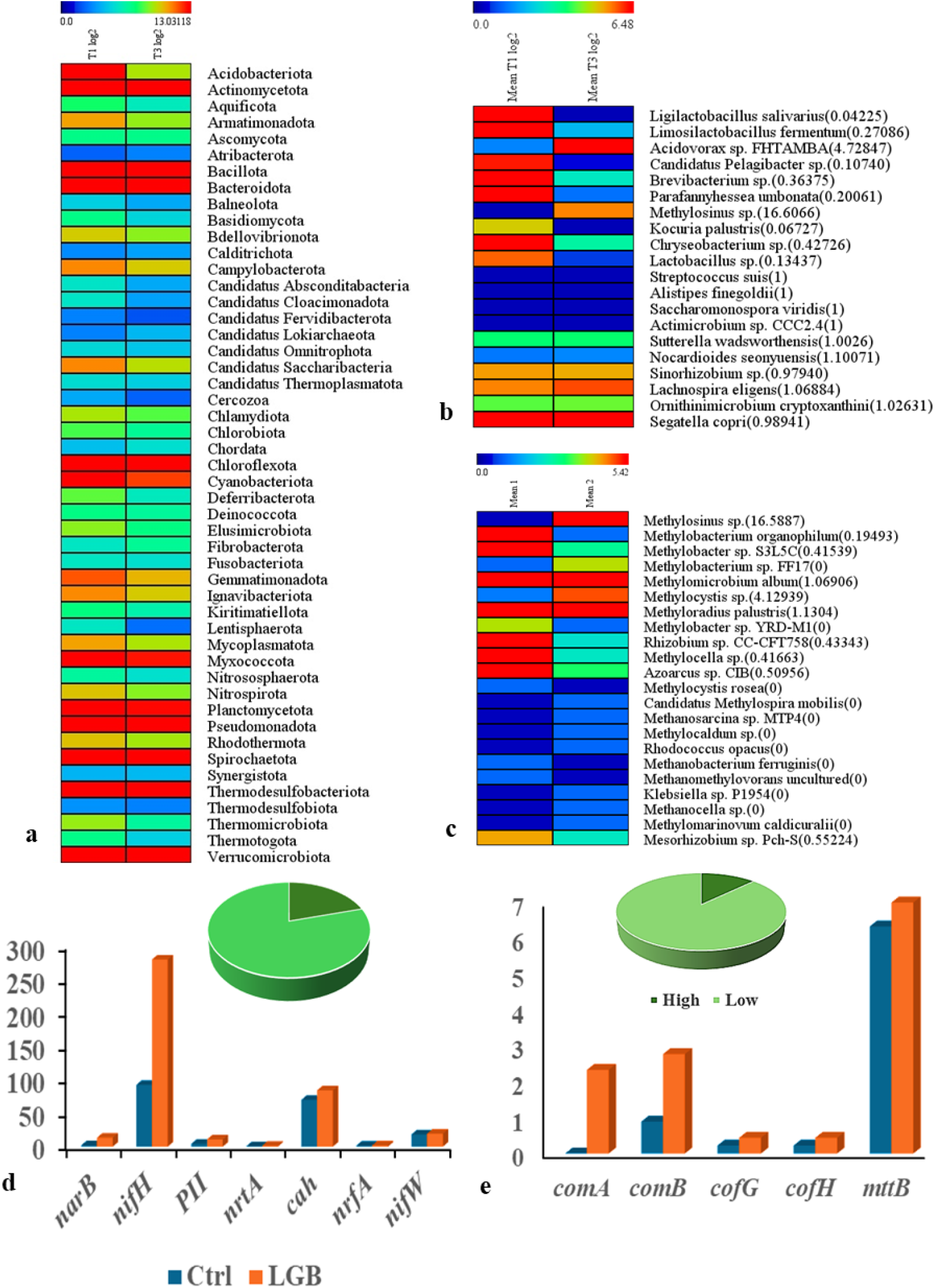
(a) Log_2_ Normalized Heatmap of control (T1) and LGB (T3) plants (inoculated GB16_1_BI) at the family level. (b) Heatmap of species with ten highest and lowest SIMPER values (values shown in parenthesis). (c) Heatmap of species predicted to be involved in N-cycling showing only ten highest and lowest SIMPER values. The values in parenthesis show the fold-change due to GB16_1_BI inoculation. (d) Analysis of nitrogen cycling genes (the pie chart depicts the manually identified OTUs) (e) Analysis of methane cycling genes (the pie chart depicts the manually identified OTUs)

Analysis of similarities (ANOSIM) values at the genus level did not reveal much difference between LGB and control plants (R= 0.2; p= 0.1). However, significant difference between sample medians of the control and LGB plants were observed by two-paired sample test and ANOVA. SIMPER analysis of normalized OTUs and fold-change analysis after log2 conversion of normalized OTUs showed that among the top ten most dissimilar OTUs, only two were higher in LGB inoculated plants (Fig. 7b). *Ligilactobacillus salivarius* had the highest dissimilarity index (0.05) but was more abundant in control plants, while *Methylosinus* sp. with the highest fold-change in inoculated samples (16-fold higher) had a dissimilarity index of 0.03. *Acidovorax* sp. FHTAMBA showed a high dissimilarity and a 4-fold higher proliferation rate in LGB plants.

Comparison of OTUs after manual functional classification (based on publicly available literature of the same species or genera), showed that OTUs with potential nitrogen-fixing ability had a significantly lower median in LGB inoculated compared to the control plants (Kruskal-Wallis test). There was a weak effect (R=0.2; p= 0.09) of LGB inoculation among the predicted N-fixing OTUs as revealed by ANOSIM analysis. *Azospirillum* sp., *Klebsiella pneumoniae, Bacillus velezensis, Azoarcus* sp., *Rhizobium* sp. and many other previously identified N-fixers were present in the samples. Among the top five most dissimilar OTUs identified from SIMPER analyses, four OTUs were repressed in LGB inoculated plants compared to control (Fig. 7c). *Methylosinus*, the most abundant OTU in LGB plants, had a high dissimilarity among the N-fixing the samples. OTUs representing the genus *Methylocystis* also showed the higher fold change in LGB plants. *Methylibium*, another methanotrophic genus capable of nitrogen-fixation was less abundant in LGB plants compared to control. Since *Methylosinus* and *Methylocystis* are known as methanotrophic genera capable of methane oxidation, the methane producing and metabolizing functional group was analysed further for diversity. However, the ANOSIM of this functional group did not show any significant influence of GB16_1_BI inoculation.

Phylogenetic investigation of communities by reconstruction of unobserved states (PICRUSt2), helped identify marker genes of functional pathways by comparing with publicly available genomes (Douglas et al., 2020). PICRUSt2 identified 4796 distinct KEGG orthologues (KO) with 618 KOs present in higher numbers in LGB plants. Among all the pathways less than 25% of the OTUs were more abundant in LGB inoculated plants. Significantly, the pentose phosphate pathway was completely repressed while 20% of the KOs representing Pentose and glucuronate interconversions were present in higher abundance in

LGB plants. A total of 38 KOs representing the nitrogen metabolism pathways were identified with 8 KOs upregulated in LGB inoculated plants. Notably, nitrogen fixation marker *nifH*, nitrate transporters and DNRA marker *nrfA* was highly abundant (**Fig 7D**). Among the methane metabolism pathway markers, 40 KOs were identified, 5 were more abundant in LGB inoculated plants. Although *comA, comB, cofG* and *cofH* are all upregulated in LGB plants the abundance is much less than the nitrogen metabolism markers (**Fig. 7E**). Another noticeable prediction from PICRUSt2 was the higher abundance of exopolysaccharide KOs in LGB plants.

## Discussion

The ability of GB16_1_BI to secrete ammonia, utilize wide range of carbohydrate, produce secondary metabolites, form persister cells and promote plant growth indicated the potential of using the strain as a bioinoculant. Formation of persister cells during nitrogen stress may help survive the harsh field conditions (Fisher et al., 2017) but makes it tougher to detect in microbiome studies due to its recalcitrant nature during DNA isolation. Persisters are slow-growing or growth-arrested cells, formed by most bacteria to survive stress conditions. Persister cells are different from the viable but non-culturable cells (Ayrapetyan et al., 2018). For assessing the feasibility of introducing GB16_1_BI as an inoculant in the field, 16S rRNA amplicon sequencing was used to study its interaction with native microbes present in the soil. Root dip inoculation of GB16_1_BI in black rice, Chakhao Poireiton, in potted soil showed that after 30 days of inoculation, majority of the microbes were supressed in LGB plants. It was also evident that GB16_1_BI encouraged the growth of selected group of microbes and promoted plant growth. Soil microbiome can be divided as the ubiquitously present generalists, and rare taxa that is specific or specialist to a soil or plant host (Pandit et al., 2009; Kim et al., 2026). While OTUs of most known nitrogen fixers were repressed in LGB plants, OTUs of microbes which are rare in soil, e.g., *Methylosinus* showed increased proliferation.

*Methylosinus* and *Methylocystis* are known to become more abundant during the seedling stages, surviving in aerobic pockets in the waterlogged soil or inside the aerenchyma tissues of rice roots (Ikeda et al., 2014; Minamisawa, 2023). It has been speculated for a long time that under the anaerobic conditions of paddy fields, methanotrophs and methanogens are important nitrogen fixers (Yoneyama et al., 2019). Similar to this study, the ability of manipulating native microbes by inoculants have been reported earlier. *Azoarcus* sp. strain KH32C encourages the proliferation of methylotrophs (Sakoda et al., 2022).

Functional prediction by PICRUSt2 showed higher *nifH, nrfA* and nitrate transport ability from the in the LGB OTUs. NifH is considered as a marker (Buckley et al., 2007) for nitrogen fixation and *nrfA* as a marker gene for DNRA (Cannon et al., 2019). While 1-aminocyclopropane-1-1carboxylate (ACC) deaminase, the marker gene for plant growth promoting bacteria, was reduced. Methane metabolism markers *comA, comB, cofG* and *cofH* were higher in LGB plant. Both *comA, comB* are required for the synthesis of Coenzyme-M and are markers for methane metabolism (Dinh and Allen et al., 2024). Similarly, *cofG* and *cofH* helps in the production of Coenzyme F420, produced by many bacterial genera (Ney et al., 2016). The cofactor F420 and coenzyme M are important in anaerobic methane production as well as oxidation (Ney et al, 2016; Dinh and Allen et al., 2024). A small-scale methane emission study done one month after transplantation in the field detected 30-60% less CH4 in LGB plants compared to control. This may be in part because GB16_1_BI can manipulate the nitrogen content, encouraging nitrogen fixers like *Methylosinus* and *Methylocystis* to proliferate. Since methanogens are known to be active from the tillering stage, it will be interesting to follow the methane production at later stages of plant growth.

Secondary metabolites produced by microbes or the host plant can also influence other microbes (Cadot et al., 2021). Hence, the ultimate consequence of plant root exudate released as a result of secondary metabolite secretion by the inoculant may lead to the reorganization of microbes over an area much larger than the 1mm^2^ soil surrounding the root (rhizosphere) (de la Porte et al., 2020). The suppression of wide range of microbes by GB16_1_BI can be correlated with the AntiMASH and metabolite data, showing its ability to produce secondary metabolites capable of influencing native microbes directly or via other keystone bacteria in the soil. A similar observation was reported after inoculation of *Streptomyces* which could interact with keystone microbes *Stenotrophomonas maltophilia* and *Paenibacillus cellulositrophicus* (Sun et al., 2025). The plant growth promotion observed in LGB plants may be either due to direct growth promoting substances produced by GB16_1_BI or due to the higher amounts of nutrients available due to suppression of native microbes.

## 5. Conclusion

In conclusion, *Microbacterium bengalense* GB16_1_BI sp. nov. (Accession number: SRX9280401) can influence a wide range of native microbe in the rhizosphere due to the production of secondary metabolites and ammonium resulting in stimulation of plant growth. Future studies may focus on different fractions of the rhizosphere separately and at different growth stages.

## Author contribution

PN: Concept, designing experiment, conducting experiment, Writing – review & editing, Writing – original draft, Funding acquisition; RG: Experimental work; KP: Experimental work; TM: Experimental work; LPC: Supervision, Writing – review & editing, resources; SD: Supervision, resources; PBMBB: Supervision and SRM: Supervision.

## Acknowledgement

PN acknowledges the Department of Science & Technology, India for funding. (SR/WOS-A/LS-463/2017 and DST/WISE/PDF/LS-4/2023). PN acknowledges the Central instrumentation and Greenhouse facility personnel of Bose Institute, Kolkata, India.

## Data availability

The WGS data is available from NCBI (SAMN48050490) and the 16S rRNA amplicon MiSeq data is available from NCBI (Bioproject ID: PRJNA1464716).

